# Human caspase-1 autoproteolysis is required for ASC-dependent and -independent inflammasome activation

**DOI:** 10.1101/681304

**Authors:** Daniel P. Ball, Cornelius Y. Taabazuing, Andrew R. Griswold, Elizabeth L. Orth, Sahana D. Rao, Ilana B. Kotliar, Darren C. Johnson, Daniel A. Bachovchin

**Affiliations:** Chemical Biology Program, Memorial Sloan Kettering Cancer Center, New York, New York 10065, USA; Weill Cornell/Rockefeller/Sloan Kettering Tri-Institutional MD-PhD Program, New York, New York 10065, USA; Tri-Institutional PhD Program in Chemical Biology, Memorial Sloan Kettering Cancer Center, New York, New York 10065, USA; Pharmacology Program of the Weill Cornell Graduate School of Medical Sciences, Memorial Sloan Kettering Cancer Center, New York, New York 10065, USA

## Abstract

Pathogen-related signals induce a number of cytosolic pattern-recognition receptors (PRRs) to form canonical inflammasomes, which activate pro-caspase-1 and trigger pyroptotic cell death. All well-studied PRRs oligomerize with the pro-caspase-1-adapter protein ASC to generate a single large structure in the cytosol, which induces the autoproteolysis and activation of the pro-caspase-1 zymogen. However, several PRRs can also directly interact with pro-caspase-1 without ASC, forming much smaller “ASC-independent” inflammasomes. It is currently thought that pro-caspase-1 autoproteolysis does not occur during, and is not required for, ASC-independent inflammasome activation. Here, we show that the related human PRRs NLRP1 and CARD8 exclusively form ASC-dependent and ASC-independent inflammasomes, respectively, identifying CARD8 as the first PRR that cannot form an ASC-containing signaling platform. Despite their different structures, we discovered that both the NLRP1 and CARD8 inflammasomes require pro-caspase-1 autoproteolysis between the small and large catalytic subunits to induce pyroptosis. Thus, pro-caspase-1 self-cleavage is an obligate regulatory step in the activation of human canonical inflammasomes.

## Introduction

Caspase-1 is a cysteine protease that induces pyroptotic cell death in response to a number of pathogen-associated signals (Broz and Dixit, 2016; Lamkanfi and Dixit, 2014). Typically, an intracellular pattern recognition receptor (PRR) senses a particular microbial structure or activity and oligomerizes with the adapter protein ASC to form an “ASC focus” in the cytosol (Broz et al., 2010a; Jones et al., 2010). The pro-caspase-1 zymogen is recruited to this structure, where it undergoes proximity-induced autoproteolysis to generate a catalytically-active enzyme. Mature caspase-1 then cleaves and activates the inflammatory cytokines pro-IL-1β and pro-IL-18 and the pore-forming protein gasdermin D (GSDMD), causing inflammatory cell death (Kayagaki et al., 2015; Shi et al., 2015). Collectively, the structures that activate pro-caspase-1 are called “canonical inflammasomes”.

Two death-fold domains, the pyrin domain (PYD) and the caspase activation and recruitment domain (CARD), mediate canonical inflammasome assembly (Broz and Dixit, 2016). ASC is comprised of a PYD and a CARD (**Fig. 1A**), and bridges either a PYD or a CARD of an activated PRR to the CARD of pro-caspase-1. In mice, all known pro-caspase-1-activating PRRs form ASC-containing inflammasomes. However, in the absence of ASC, two murine CARD-containing PRRs, NLRC4 and NLRP1B, can directly recruit and activate pro-caspase-1 through CARD-CARD interactions (Broz et al., 2010b; Guey et al., 2014; Mariathasan et al., 2004; Poyet et al., 2001; Van Opdenbosch et al., 2014). ASC-independent inflammasomes induce the cleavage of GSDMD and trigger lytic cell death, but do not form large foci or efficiently process pro-caspase-1 and pro-IL-1β (Broz et al., 2010b; He et al., 2015).

**Figure 1.**
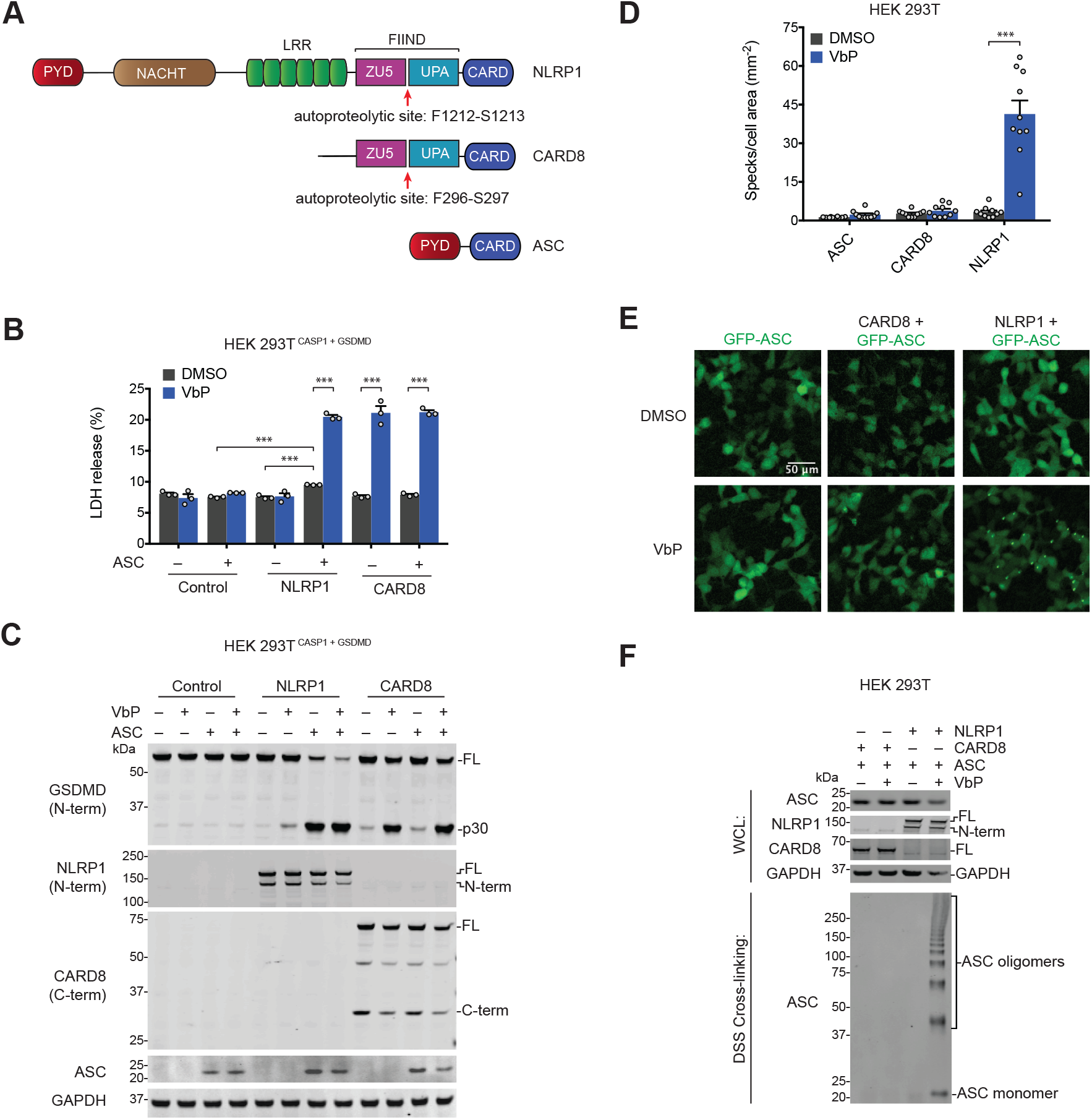
NLRP1 is ASC-dependent and CARD8 is ASC-independent. **(A)** Schematic for human NLRP1, CARD8, and ASC protein domain structures. The autoproteolysis sites are indicated. The ZU5-UPA domains together are also referred to as a FIIND. **(B,C)** HEK 293T cells stably expressing CASP1 and GSDMD (HEK 293T^CASP1 + GSDMD^) were transfected with constructs encoding the indicated proteins and treated with DMSO or VbP (10 μM, 6 h). Supernatants were evaluated for LDH release **(B)** and lysates were analyzed by immunoblotting **(C)**. Data are means ± SEM of three biological replicates. *** *p* < 0.001 by two-sided Students *t*-test. FL, full-length. **(D,E)** HEK 293T cells were transfected with constructs encoding GFP-tagged ASC and NLRP1 or CARD8, treated with DMSO or VbP (10 μM, 6 h), and evaluated for ASC speck formation by fluorescence microscopy. Shown are the mean ± SEM **(D)** and representative images **(E)** from 10 technical replicates from one of two independent experiments. *** *p* < 0.001 by two-sided Students *t*-test. **(F)** HEK 293T cells transiently transfected with constructs encoding the indicated proteins and treated with DMSO or VbP (10 μM, 6 h). Lysates were harvested, subjected to DSS crosslinking, and evaluated by immunoblotting.

These observations suggested that pro-caspase-1 autoproteolysis may not be required for cell death. To explore this possibility, two independent groups reconstituted *Casp1^−/−^* mouse macrophages (which expressed ASC) with an uncleavable mutant form of mouse pro-caspase-1, and found that the mutant enzyme still mediated cell death, but did not process pro-IL-1β, in response to various inflammasome stimuli (Broz et al., 2010b; Guey et al., 2014). Another study, performed after the discovery of GSDMD, showed that the uncleavable mutant pro-caspase-1 was partially defective in processing GSDMD and inducing pyroptosis in ASC-expressing RAW 264.7 cells in response to NLRP3 inflammasome activation (He et al., 2015). Taken together, these studies indicated that murine ASC-containing inflammasomes can activate pro-caspase-1 to some extent, but that autoproteolysis was required for full catalytic activity. Because ASC-independent inflammasomes induce little detectable pro-caspase-1 and pro-IL-1β processing, it has been widely assumed that ASC-independent inflammasomes specifically activate the pro-protein form of caspase-1 without autoproteolysis. However, the importance of pro-caspase-1 autoproteolysis in mouse ASC-independent inflammasome activation has not been directly tested, perhaps in part because these structures are not known to form in physiologically-relevant macrophages that express ASC. Moreover, the requirement of human pro-caspase-1 autoproteolysis in the activation of either ASC-independent or ASC-dependent inflammasomes has not been evaluated experimentally.

DPP8/9 inhibitors activate the related CARD-containing human NLRP1 and CARD8 inflammasomes (**Fig. 1A**), which both have C-terminal ZU5, UPA, and CARD domains (Chui et al., 2019; Johnson et al., 2018; Okondo et al., 2017; Zhong et al., 2018). The ZU5 domains of NLRP1 and CARD8 undergo post-translational autoproteolysis (**Fig. 1A**), generating non-covalently associated, autoinhibited N- and C-terminal polypeptide fragments (D’Osualdo et al., 2011; Finger et al., 2012; Frew et al., 2012). The C-terminal UPA-CARD fragments mediate cell death (Finger et al., 2012; Johnson et al., 2018). CARD8 does not require ASC to activate pro-caspase-1 (Johnson et al., 2018; Okondo et al., 2017), but it is unknown whether CARD8 can also form an ASC-containing inflammasome. In contrast, human NLRP1, unlike mouse NLRP1A and NLRP1B (Masters et al., 2012; Van Opdenbosch et al., 2014), appears to require ASC (Finger et al., 2012; Zhong et al., 2016; Zhong et al., 2018). Here, we show that CARD8 and NLRP1 exclusively form ASC-independent and ASC-dependent inflammasomes, respectively, due to specific CARD-CARD interactions. These data identify CARD8 as the first pro-caspase-1-activating PRR that cannot form an ASC focus. Although the CARD8 inflammasome induces little detectable pro-caspase-1 processing by immunoblotting (Johnson et al., 2018; Okondo et al., 2017), we found that pro-caspase-1 autoproteolysis was required for activation of both the CARD8 and NLRP1 inflammasomes. Overall, these data demonstrate that autoproteolysis is critical for the activation of human canonical inflammasomes.

## Results and discussion

### NLRP1 is ASC-dependent and CARD8 is ASC-independent

We first wanted to determine the capabilities of human NLRP1 and CARD8 to form ASC-dependent and ASC-independent inflammasomes. We therefore transfected constructs encoding NLRP1, CARD8, and/or ASC into HEK 293T cells stably expressing pro-caspase-1 and GSDMD before treatment with the DPP8/9 inhibitor Val-boroPro (VbP). VbP induced similar levels of GSDMD cleavage and LDH release in cells expressing CARD8 in the presence or absence of ASC (**Fig. 1B,C**), confirming that ASC is not required for CARD8-mediated cell death (Johnson et al., 2018; Okondo et al., 2017). In contrast, NLRP1 required ASC co-expression to mediate cell death (**Fig. 1B,C**). We should note that the co-expression of NLRP1 and ASC induced some spontaneous cell death and GSDMD cleavage, but both were increased by VbP. Consistent with these data, transient transfection of constructs encoding the active UPA-CARD fragment of NLRP1, but not CARD8, required ASC to induce GSDMD cleavage (**Fig. S1A**). As previously reported, the PYD of NLRP1 was dispensable for inflammasome activation (**Fig. S1B,C**) (Chavarria-Smith et al., 2016; Finger et al., 2012).

Although these results confirm that CARD8 can directly activate pro-caspase-1 without ASC bridging, it remained possible that CARD8 could also form an ASC-containing inflammasome, similar to mouse NLRP1B (Van Opdenbosch et al., 2014). We next co-transfected HEK 293T cells with constructs encoding GFP-tagged ASC and either NLRP1 or CARD8. These cells were then treated with VbP for 6 h and imaged by fluorescence microscopy (**Fig. 1D,E**). VbP induced ASC specks in NLRP1, but not CARD8, expressing cells, suggesting that CARD8 cannot form an ASC-containing inflammasome. Similarly, transfection of the UPA-CARD of NLRP1, but not CARD8, induced ASC speck formation (**Fig. S1D,E**). To further support these microscopy results, we co-transfected HEK 293T cells with constructs encoding untagged ASC and either NLRP1 or CARD8, treated the cells with VbP, and cross-linked lysates with disuccinimidyl suberate (DSS). As expected, VbP induced ASC oligomerization in cells expressing NLRP1, but not CARD8 (**Fig. 1F**).

We hypothesized that the exclusive formation of ASC-independent and ASC-dependent inflammasomes by CARD8 and NLRP1, respectively, was due to specific interaction differences between the CARDs of CARD8 and NLRP1 with the CARDs of ASC and CASP1. To test this prediction, we incorporated these CARDs into a split luciferase-based NanoBiT assay (Dixon et al., 2016), fusing Small BiT (SmBiT, an 11 amino acid peptide) to the CARD domains of ASC and CASP1 and Large BiT (LgBiT, an 18 kDa tag that luminesces only when bound to SmBiT) to the CARD domains of ASC, CASP1, CARD8, and NLRP1 (**Fig. 2A**). We mixed lysates containing the indicated fusion proteins, and observed luminescent signals indicating binding between the ASC^CARD^ and itself, CASP1^CARD^, and NLRP1^CARD^ (**Fig. 2B**), and between the CASP1^CARD^ and itself, ASC^CARD^, and CARD8^CARD^ (**Fig. 2C**). As expected, we did not observe a CASP1^CARD^-NLRP1^CARD^ interaction or an ASC^CARD^-CARD8^CARD^ interaction. Overall, these results indicate specific CARD-CARD interactions govern the formation of the CARD8 ASC-independent inflammasome and the NLRP1 ASC-dependent inflammasome.

**Figure 2.**
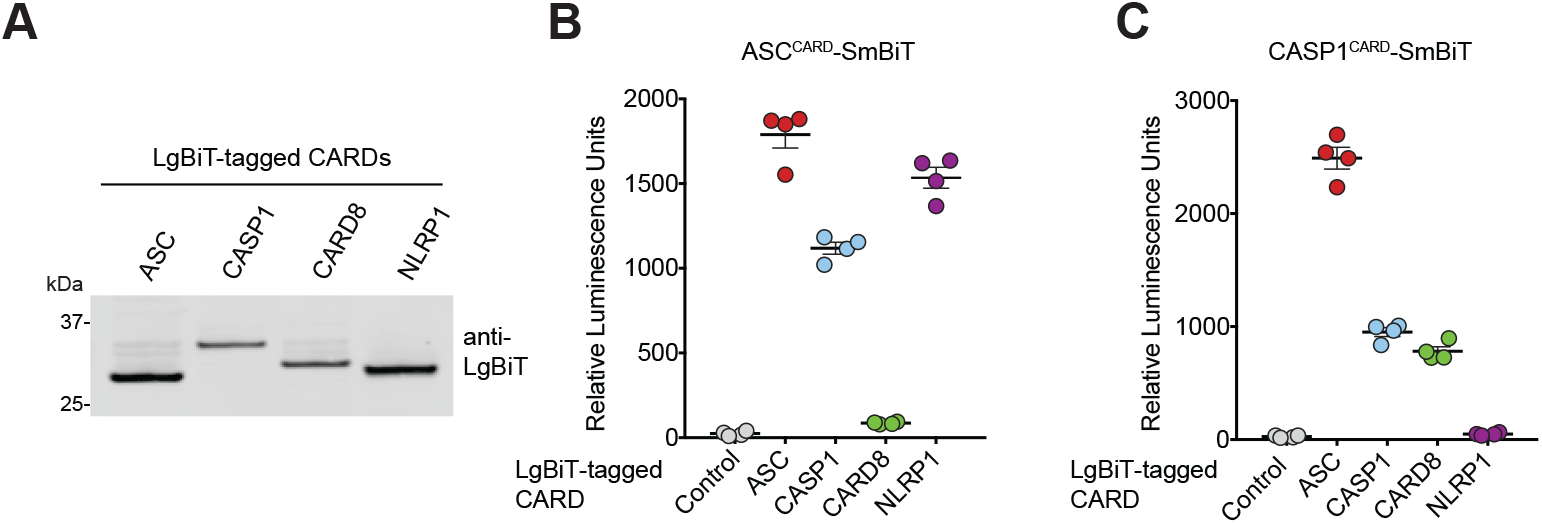
Specific CARD-CARD interactions determine ASC-dependent or independent inflammasome assembly. **(A)** Expression of the indicated LgBiT-tagged CARDs in HEK 293T cells was verified by immunoblotting. **(B,C)** Cell lysates from HEK 293T cells transiently expressing LgBiT-tagged ASC^CARD^ **(B)** or LgBiT-tagged CASP1^CARD^ **(C)** were mixed with lysates containing SmBiT-tagged CARDs and analyzed for the relative luminescence.

### Proteasome activity is critical for NLRP1 activation

DPP8/9 inhibition induces the proteasome-mediated degradation of the N-terminal fragment of mouse NLRP1B and CARD8, releasing the UPA-CARD C-terminal fragment to activate pro-caspase-1 (Chui et al., 2019; Johnson et al., 2018). Given the differences in the N-terminal regions of CARD8 and NLRP1 (**Fig. 1A**), we wanted to confirm that VbP activates human NLRP1 by a similar degradation mechanism. Indeed, autoproteolysis-defective NLRP1 S1213A, which is unable to release its C-terminal fragment, was severely impaired in VbP-induced ASC speck formation (**Fig. S2A,B**) and cell death (**Fig. S2C,D**). Moreover, proteasome inhibitors partially rescued VbP-induced cell death (**Fig. 3A, Fig. S1B**), GSDMD cleavage (**Fig. 3B, Fig. S1C**), and ASC oligomerization (**Fig. 3C**). We speculate that proteasome blockade did not fully rescue VbP-induced NLRP1 activation because very small amounts of UPA-CARD are needed to nucleate ASC specks (Sandstrom et al., 2019), and therefore even slight residual proteasome activity could be sufficient to activate the inflammasome. Consistent with only a small amount of NLRP1 UPA-CARD being liberated, VbP did not induce obvious NLRP1 protein depletion by immunoblotting in several experiments (**Fig. 1C, Fig. 3B**). However, we confirmed that VbP does indeed induce NLRP1 protein depletion by treating NLRP1-expressing HEK 293T cells with VbP for longer time periods (**Fig. S2E**). It should be noted that these cells do not express pro-caspase-1, and thus NLRP1 loss here is not due to selective elimination of NLRP1-expressing cells.

**Figure 3.**
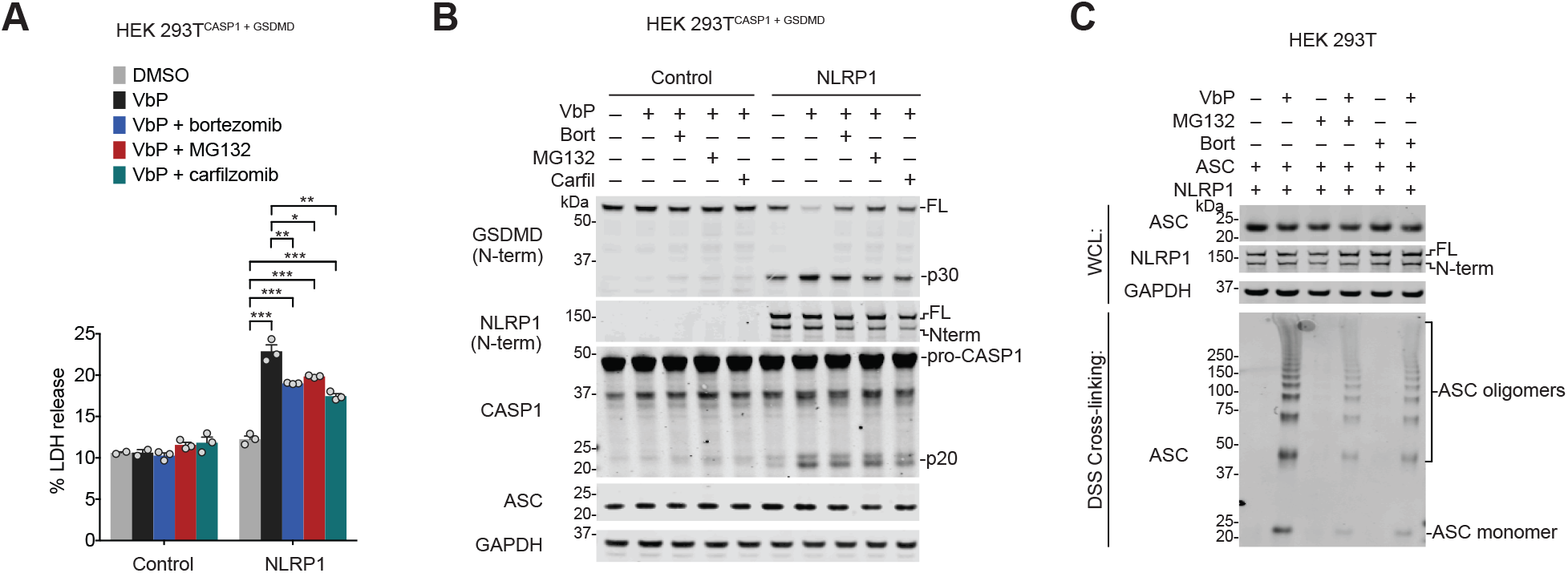
Proteasome inhibitors block NLRP1 inflammasome activation. **(A,B)** HEK 293T^CASP1 + GSDMD^ were transiently transfected with constructs encoding NLRP1 and ASC, pretreated with the indicated proteasome inhibitors (20 μM, 30 min), and stimulated with VbP (10 μM, 6h). Supernatants were evaluated for LDH release **(B)** and lysates were analyzed by immunoblotting **(C)**. Data are means ± SEM of three biological replicates and representative of two independent experiments. * *p* < 0.05*, * *p* < 0.01, *** *p* < 0.001 by two-sided Students *t*-test. **(C)** HEK 293T cells transiently transfected with constructs encoding NLRP1 and ASC, preincubated with MG132, carfilzomib, or bortezomib (20 μM, 30 min), and treated with DMSO or VbP (10 μM, 6 h).

Germline mutations in the N-terminal fragment of NLRP1 cause several related inflammatory skin disorders (Zhong et al., 2016; Zhong et al., 2018). We hypothesized that these mutations destabilized the N-terminal fragment, leading to increased proteasome-mediated N-terminal degradation. Indeed, we found that the proteasome inhibitor bortezomib reduced spontaneous inflammasome activation caused by several of these mutations (**Fig. S2F,G**). Overall, these data indicate that the proteasome mediates both VbP- and mutation-induced NLRP1 activation.

### The ASC-independent inflammasome requires caspase-1 processing

We next wanted to study the requirements for ASC-independent inflammasome activation in greater detail. We initially discovered DPP8/9 inhibitor-induced pyroptosis in human THP-1 cells (Okondo et al., 2017), which is mediated by CARD8 (Johnson et al., 2018). We observed little, if any, caspase-1 and IL-1β processing, and designated this death as “pro-caspase-1 dependent” pyroptosis. However, as described above, we never formally demonstrated that pro-caspase-1 itself mediates this response. We reasoned that we might observe more caspase-1 processing, if it was occurring, in *GSDMD^−/−^* THP-1 cells, as the cleaved products would not be as readily released into the supernatant. Indeed, we did observe bands corresponding to the p35 and p20 fragments in these knockout cells (**Fig. 4A-C**), indicating that the CARD8 inflammasome can, in fact, process pro-caspase-1. It should be noted that VbP induces apoptosis in *GSDMD^−/−^* THP-1 cells (Taabazuing et al., 2017), and as expected PARP cleavage was observed here.

**Figure 4.**
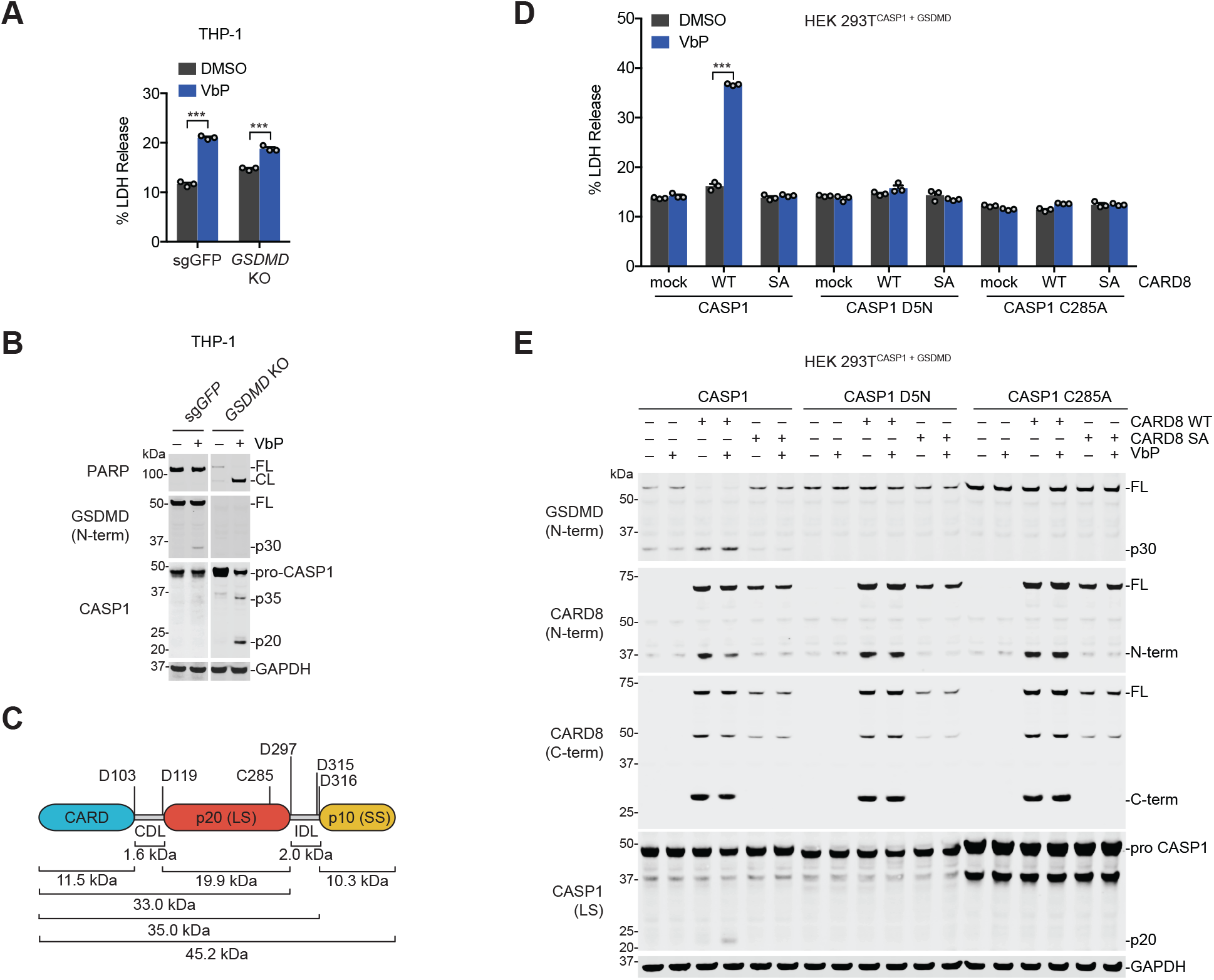
Caspase-1 autoproteolysis is required for CARD8 inflammasome activation. **(A,B)** Control and *GSDMD^−/−^* THP-1 cells were treated with VbP (10 μM, 24 h) before supernatants were analyzed for LDH release **(A)** and lysates were evaluated by immunoblotting **(B)**. Data are means ± SEM of three biological replicates. *** *p* < 0.001 by two-sided Students *t*-test. FL, full-length. CL, cleaved. **(C)** Schematic of pro-caspase-1 depicting the CARD domain and large (p20, LS) and small (p10, SS) catalytic subunits. Predicted cleavage sites, sizes of potential cleavage products, and the catalytic cysteine are indicated. **(D, E)** HEK 293T cells stably expressing GSDMD and the indicated pro-caspase-1 constructs were transiently transfected with plasmids encoding RFP (mock), CARD8 WT, or autoproteolysis-defective CARD8 S297A (SA) for 24 h before addition of VbP (10 μM, 6 h). Cell death was assessed by LDH release **(D)** and GSDMD and CASP1 cleavage by immunoblotting **(E)**. Data are means ± SEM of three biological replicates. *** *p* < 0.001 by two-sided Students *t*-test.

We next wanted to determine if caspase-1 processing was required for cell death. Analogous to the previously created uncleavable mouse pro-caspase-1 (mCASP1 D6N) (Broz et al., 2010b), we generated an uncleavable human pro-caspase-1 (CASP1 D5N, **Fig. 4C**) in which all Asp cleavage sites were mutated to Asn residues (Thornberry et al., 1992). We then created HEK 293T cell lines stably expressing wild-type (WT), uncleavable (D5N), or catalytically-inactive (C285A) pro-caspase-1, transiently transfected constructs encoding WT or autoproteolytic-defective S297A CARD8 into each these cell lines, and treated with VbP. As expected, we observed robust cell death and GSDMD cleavage in cells with WT pro-caspase-1 and WT CARD8, but not in cells expressing catalytically-dead CASP1 or autoproteolysis-defective CARD8 (**Fig. 4D**). Interestingly, we also observed a small amount of the p20 cleaved product in the cell line expressing CASP1 WT. In contrast, we did not observe any cell death or GSDMD cleavage in cells expressing the uncleavable CASP1 D5N. Consistent with these data, transient transfection of a plasmid encoding the active UPA-CARD fragment of CARD8 did not induce GSDMD cleavage in cells expressing CASP1 D5N (**Fig. S3A**). Together, these data indicate that pro-caspase-1 autoproteolysis is required for CARD8 inflammasome activation.

### Cleavage in the caspase-1 interdomain linker (IDL) is essential for activation

We next wanted to determine which specific pro-caspase-1 cleavage sites were required for CARD8 inflammasome activation, and to determine if pro-caspase-1 autoproteolysis was also required for NLRP1 inflammasome activation. Pro-caspase-1 is comprised of three domains, a CARD, a large subunit (LS, p20), and a small subunit (SS, P10), separated by two linkers (**Fig. 4C**). Pro-caspase-1 undergoes proteolytic processing at two sites (D103 and D119) in the CARD linker (CDL) that separates the CARD and the p20, and three sites (D297, D315, and D316) in the interdomain linker (IDL) that separates the p20 and the p10 (Boucher et al., 2018; Thornberry et al., 1992). As IDL cleavage has been associated with higher catalytic activity and CDL cleavage with termination of activity (Boucher et al., 2018; Broz et al., 2010b), we first tested the 3 putative cleavage sites in the IDL by generating HEK 293T cells stably expressing CASP1 D297N, D315N/D316N, and D297N/D315N/D316N (“IDL uncleavable”, or IDL^uncl^). We then transfected plasmids encoding CARD8 or both NLRP1 and ASC into these cell lines and treated with VbP. We found that VbP induced LDH release and GSDMD cleavage in cells expressing CASP1 D297N and CASP1 D315N/D315N, but not in cells expressing CASP1 IDL^uncl^ (**Fig. 5**). These data show that pro-caspase-1 autoproteolysis within the IDL is critical for both ASC-independent and -dependent inflammasome activation. As predicted by these results, transfection of plasmids encoding ASC, the UPA-CARD of CARD8, or residues 1-328 of human NLRC4, which contains a CARD domain that can directly activate human CASP1 (Poyet et al., 2001), failed to induce death in the cells expressing CASP1 IDL^uncl^ (**Fig. S3B-D**). In contrast to human CASP1 D5N and consistent with previous reports (Broz et al., 2010b; Guey et al., 2014), the UPA-CARD of mouse NLRP1B induced cell death in HEK 293T cells stably expressing the mouse CASP1 D6N protein (**Fig. S3E,F**). Surprisingly, however, the NLRP1B UPA-CARD also induced the formation of several lower molecular weight caspase-1 species in the CASP1 D6N-expressing line, indicating that the mouse CASP1 D6N protein is, in fact, cleavable. These potential additional cleavage sites and their function in mouse caspase-1 activation warrant future investigations.

**Figure 5.**
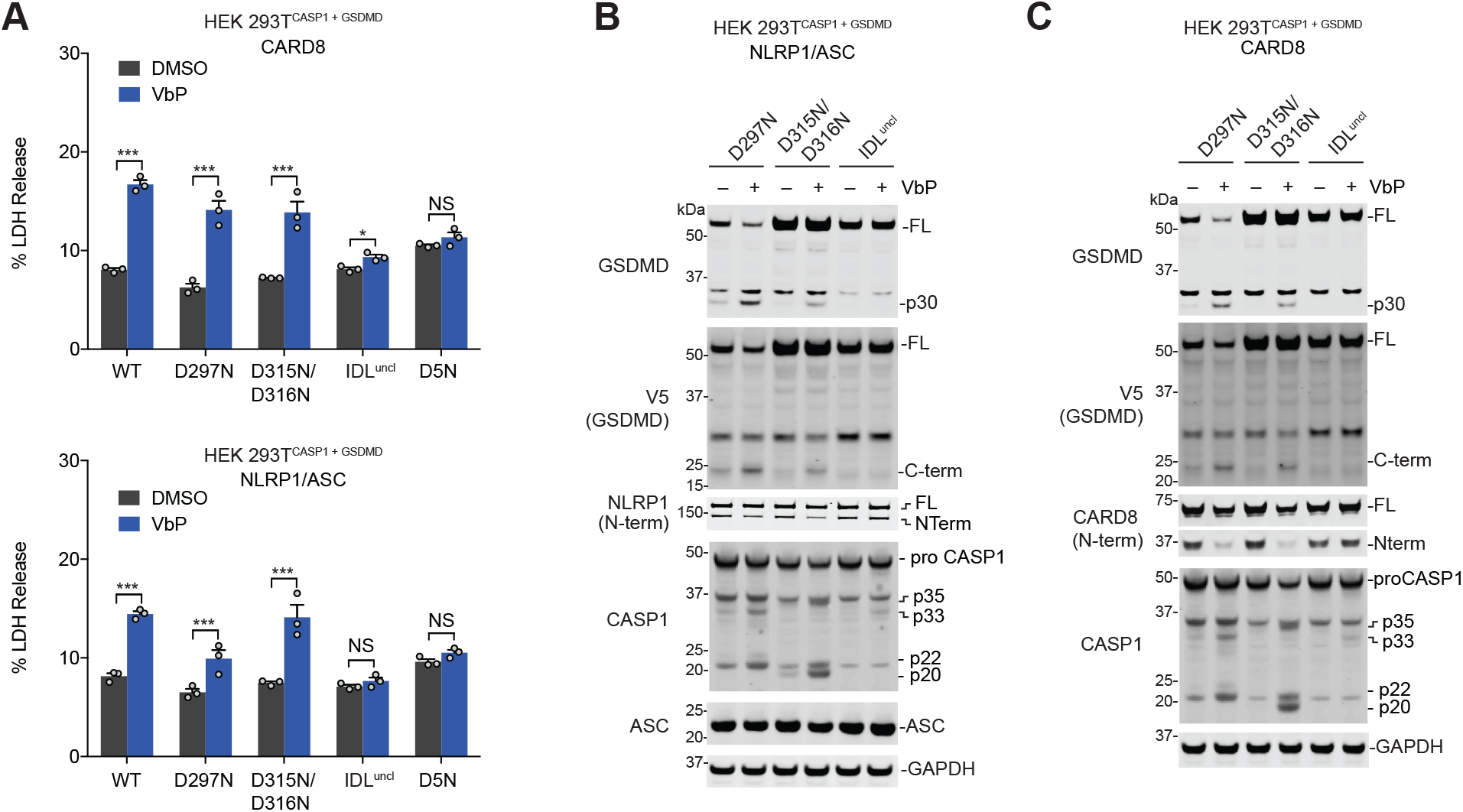
Cleavage of the human caspase-1 IDL is required for activation of canonical inflammasomes. **(A-C)** HEK 293T cells stably expressing GSDMD and the indicated pro-caspase-1 constructs were transiently transfected with plasmids encoding NLRP1 (0.1 μg) and ASC (0.01 μg) **(A,B)** or CARD8 **(A,C)** for 24 h before addition of VbP (10 μM, 6 h). Cell death was assessed by LDH release **(A)** and GSDMD cleavage by immunoblotting **(B,C)**. Data are means ± SEM of three biological replicates. * *p* < 0.05, *** *p* < 0.001 by two-sided Students *t*-test. NS, not significant.

Replacing CARDs with DmrB domains enables the small-molecule (AP-20187)-induced dimerization and activation of caspases (Boucher et al., 2018; Ross et al., 2018; Ruhl et al., 2018). To confirm that IDL cleavage was required for proximity-induced pro-caspase-1 activation, we cloned DmrB-caspase-1 constructs with the IDL mutations described above (**Fig. S3G**). We transiently transfected these constructs into HEK 293T cells, and then treated these cells with the AP-20187. We observed that the WT DmrB-caspase-1 underwent significant autoproteolysis and triggered GSDMD cleavage (**Fig. S3H**). Some pro-caspase-1 autoproteolysis and GSDMD cleavage were also observed for D297N and the D315N/D316N mutants, but not the IDL^uncl^ mutant. Thus, these data confirm the importance of IDL processing for human caspase-1 activation.

Here, we have shown that the related NLRP1 and CARD8 inflammasomes are remarkably distinct. First, these PRRs have functionally divergent C-terminal UPA-CARD fragments – one that induces an ASC focus to indirectly activate pro-caspase-1 and one that directly activates pro-caspase-1. As such, we predict that the physiological outputs of NLRP1 and CARD8 activation will be different *in vivo*, for example in the kinetics of immune activation or in the type or extent of cytokine processing. Future investigations are needed to establish the biological purpose of ASC-independent and ASC-dependent inflammasomes. Second, CARD8 and NLRP1 have entirely dissimilar N-terminal fragments. Although both are activated by at least one similar signal—the cellular consequence of DPP8/9 inhibition—we speculate that these N-terminal fragments likely evolved for different purposes that remain to be elucidated.

More generally, we have now demonstrated that human pro-caspase-1 autoproteolysis is necessary for both ASC-dependent and ASC-independent inflammasome activation. Interestingly, two recent studies have demonstrated that the related inflammatory caspase-11, which only forms an ASC-independent inflammasome (termed the “non-canonical” inflammasome) with often little detectable self-cleavage and no IL-1β processing (Hagar et al., 2013; Yang et al., 2015), also requires IDL autoproteolysis for activation (Boucher et al., 2018; Lee et al., 2018). In this way, the ASC-independent caspase-1 canonical inflammasome is remarkably similar to the non-canonical caspase-11 inflammasome. Collectively, these reports and our data show that limited proteolysis plays a critical role in the activation of inflammatory caspases.

## Materials and Methods

### Antibodies and reagents

Antibodies used include: GSDMD Rabbit polyclonal Ab (Novus Biologicals, NBP2-33422), human NLRP1/NALP1 Sheep polyclonal antibody (R&D systems, AF6788), V5 Rabbit polyclonal Ab (Abcam, Ab9116), FLAG^®^ M2 monoclonal Ab (Sigma, F3165), CARD8 N-terminus Rabbit polyclonal antibody (Abcam, Ab194585), CARD8 C-terminus Rabbit polyclonal Ab (Abcam, Ab24186), human ASC Sheep polyclonal antibody (R&D systems, AF3805), GAPDH Rabbit monoclonal Ab (Cell Signaling Tech, 14C10), NLuc (Lg-BiT) polyclonal antibody (courtesy of Promega), human Caspase-1 p20 Rabbit polyclonal Ab (Cell Signaling Technology, #2225), PARP Rabbit polyclonal Ab (Cell Signaling Technology, #9542), IRDye 680 RD Streptavidin, (LI-COR 926-68079), IRDye 800CW anti-rabbit (LICOR, 925-32211), IRDye 800CW anti-mouse (LI-COR, 925-32210), IRDye 680CW anti-rabbit (LI-COR, 925-68073), IRDye 680CW anti-mouse (LI-COR, 925-68072). Other reagents used include: Val-boroPro (VbP)(Okondo et al., 2017), Bortezomib (MilliporeSigma, 504314), MG132 (MilliporeSigma, 474790), Carfilzomib (Cayman Chemical, 17554), B/B Homodimerizer (Takara, 635059, equivalent to AP-20187), disuccinimidyl suberate (DSS, ThermoFisher Scientific, 21655), FuGENE HD (Promega, E2311).

### Cell Culture

HEK 293T cells and THP-1 cells were purchased from ATCC. HEK 293T cells were grown in Dulbecco’s Modified Eagle’s Medium (DMEM) with L-glutamine and 10% fetal bovine serum (FBS). THP-1 cells were grown in Roswell Park Memorial Institute (RPMI) medium 1640 with L-glutamine and 10% fetal bovine serum (FBS). All cells were grown at 37 °C in a 5% CO_2_ atmosphere incubator. Cell lines were regularly tested for mycoplasma using the MycoAlert™ Mycoplasma Detection Kit (Lonza). Stable cell lines were generated as described previously (Johnson et al., 2018).

### Cloning

Plasmids for full-length and truncated CARD8, NLRP1, mouse NLRP1B (allele 1), mouse and human GSDMD, and mouse and human CASP1 (Johnson et al., 2018; Okondo et al.; Okondo et al., 2018) were cloned as previously described and shuttled into modified pLEX_307 vectors (Addgene) using Gateway technology (Thermo Fisher Scientific). A plasmid encoding NLRC4 was purchased from Origene (RC206757) and cloned into the Gateway system. A pLEX_307 vector containing RFP was used for controls. Point mutations were generated using the QuickChange II site-directed mutagenesis kit (Agilent, 200523) following the manufacturer’s instructions. The NLRP1ΔPYD construct starts at Ser93. DNA encoding SmBit and LgBit for the NanoBiT assay (Promega) were inserted after the attR2 recombination site in a modified pLEX_307 vector (immediately after the EcoRV site), and DNA encoding CARD domains were shuttled into these modified vectors using Gateway technology. DmrBΔCARD caspase-1 chimera constructs were cloned using assembly PCR reactions beginning at Asp92 of caspase-1.

### Transient transfections

HEK 293T cells were plated in 6-well culture plates at 5.0 × 10^5^ cells/well in DMEM. The next day, the indicated plasmids were mixed with an empty vector to a total of 2.0 μg DNA in 125 μl in Opti-MEM and transfected using FuGENE HD (Promega) according to the manufacturer’s protocol. Unless indicated otherwise, 0.02 μg CARD8, 0.02 μg NLRP1, and 0.005 μg ASC were used. The next day, the cells were treated as described. For microscopy experiments, cells were plated directly into Nunc Lab-Tek II Chamber slide w/Cover sterile glass slides (Thermo Fisher Scientific, 154534) at 8.0 × 10^4^ cells/well and treated with 25 μL transfection master mix dropwise.

### LDH cytotoxicity and immunoblotting assays

HEK 293T cells were transiently transfected and inhibitor treated as indicated. THP-1 cells were plated in 6-well culture plates at 5.0 × 10^5^ cells/well and treated with VbP as indicated. 15 min prior to the conclusion of cell transfection experiments 80 μL of a 9% Triton X-100 solution was added to designated lysis control wells of a 6-well culture plate to completely lyse the cell contents. Supernatants were analyzed for LDH activity using the Pierce LDH Cytotoxicity Assay Kit (Life Technologies) and lysates protein content was evaluated by immunoblotting. Cells were washed 2× in PBS (pH = 7.4), resuspended in PBS, and lysed by sonication. Protein concentrations were determined using the DCA Protein Assay kit (Bio-Rad). The samples were separated by SDS-PAGE, immunoblotted, and visualized using the Odyssey Imaging System (Li-Cor).

### Fluorescence microscopy

Imaging was performed on a Zeiss Axio Observer.Z1 inverted widefield microscope using 40x/0.95NA air objective. Cells were plated on LabTek 8-well chambered cover glass with #1 coverslip. For each chamber, 10 positions were imaged with brightfield, red, and green fluorescence channels as a single time point at the conclusion of the given experiment. Data was exported as raw .czi files and analyzed using custom macro written in ImageJ/FIJI. Total cell area was estimated from RFP-positive signal and the number of GFP-ASC specks were quantified using the “Analyze particles” function following threshold adjustment in the GFP positive images.

### Split luciferase assay

HEK cells were seeded at 3.0 × 10^6^ cells in 10 cm dishes and transfected with 3 μg of the indicated DNA construct using FuGENE HD (Promega). 24 h post-transfection, cells were washed with cold PBS (Corning), harvested by scraping, and pelleted at 450 × *g* for 3 min. The pellets were resuspended in 500 μL PBS and lysed by sonication. Lysates were clarified to remove bulk cellular debris by centrifugation at 1000 × *g* for 5 min, and relative expression was normalized by gel densitometry of immunoblots (ImageJ 1.52n software). NanoBiT assays were carried out in quadruplicate in white, clear, flat-bottom, 384-well assay plates (Corning, 3765). Equal volume aliquots of the corresponding SmBiT/LgBiT pairs were combined within each well from normalized lysates, followed by addition of Nano-Glo Live Cell Reagent, prepared as per manufacturer’s instructions. Following thermal equilibration, luminescence was read on a Cytation 5 multi-modal plate reader.

### DSS Cross-linking

HEK 293T cells were treated as indicated before lysates were harvested and pelleted at 400 × g 4 °C for 3 min and washed with cold PBS. Cell pellets were lysed with 200 μL 0.5% NP-40 in TBS for 30 min on ice in 1.75 mL microcentrifuge tubes. The lysates were spun down at 1,000 g 4 °C for 10 min to remove bulk cell debris (100 μL of supernatant was reserved for immunoblot). The remaining lysate was placed in the centrifuge for 10 min at 20,000 x *g* 4 °C. The obtained pellet was then washed with 100 μL CHAPS buffer (50 mM HEPES pH 7.5, 5 mM MgCl2, 0.5 mM EGTA, and 0.1% w/v CHAPS) then resuspended in 48 μL CHAPS buffer. 2 μL of 250 mM DSS was added and the samples were agitated at 37 °C on a rotating orbital platform set to 1,000 rpm for 45 min to facilitate protein cross-linking. The samples were then combined with an equal volume of 2× loading dye and heated to 98 °C for 10 min and prepared for immunoblot analysis.

### Data analysis and statistics

Statistical analysis was performed using GraphPad Prism 7.0 software. Statistical significance was determined using two-sided Students *t*-tests.

## Supplemental material

Fig. S1 shows additional data related to Fig. 1 and Fig. 2, demonstrating that the UPA-CARD of NLRP1, but not CARD8, interacts with ASC and requires ASC to activate pro-caspase-1. Fig. S2 displays additional data related to Fig. 3, confirming that the proteasome plays an important role in the activation of human NLRP1. In particular, this figure shows NLRP1 autoproteolysis is required, that VbP induces NLRP1 protein loss, and spontaneous activation of NLRP1 by germline mutations is blocked by bortezomib. Fig. S3 shows data related to Fig. 4 and Fig 5., confirming that human caspase-1 autoproteolysis within the IDL is required for inflammasome activation.

## Author Contributions

D.P.B, C.Y.T., A.R.G, S.D.R., and D.C.J. performed experiments. D.P.B, C.Y.T., and D.A.B. designed experiments, analyzed data, and wrote the paper. I.B.K. and E.L.O. developed and performed the split luciferase assay.

## Acknowledgements

We thank K. Schroder and P. Broz for sharing DmrB constructs. This work was supported by the Josie Robertson Foundation (D.A.B.), a Stand Up to Cancer-Innovative Research Grant (Grant Number SU2C-AACR-IRG11-17 to D.A.B.; Stand Up to Cancer is a program of the Entertainment Industry Foundation. Research Grants are administered by the American Association for Cancer Research, the scientific partner of SU2C), the Pew Charitable Trusts (D.A.B. is a Pew-Stewart Scholar in Cancer Research), the Pershing Square Sohn Cancer Research Alliance (D.A.B.), the NIH (R01 AI137168 to D.A.B.; T32 GM007739-Andersen to A.R.G; the MSKCC Core Grant P30 CA008748), an Alfred P. Sloan Foundation Research Fellowship (D.A.B.), Gabrielle’s Angel Foundation (D.A.B.), and the American Cancer Society (Postdoctoral Fellowship PF-17-224-01 – CCG to C.Y.T.).

---

**Figure S1.**
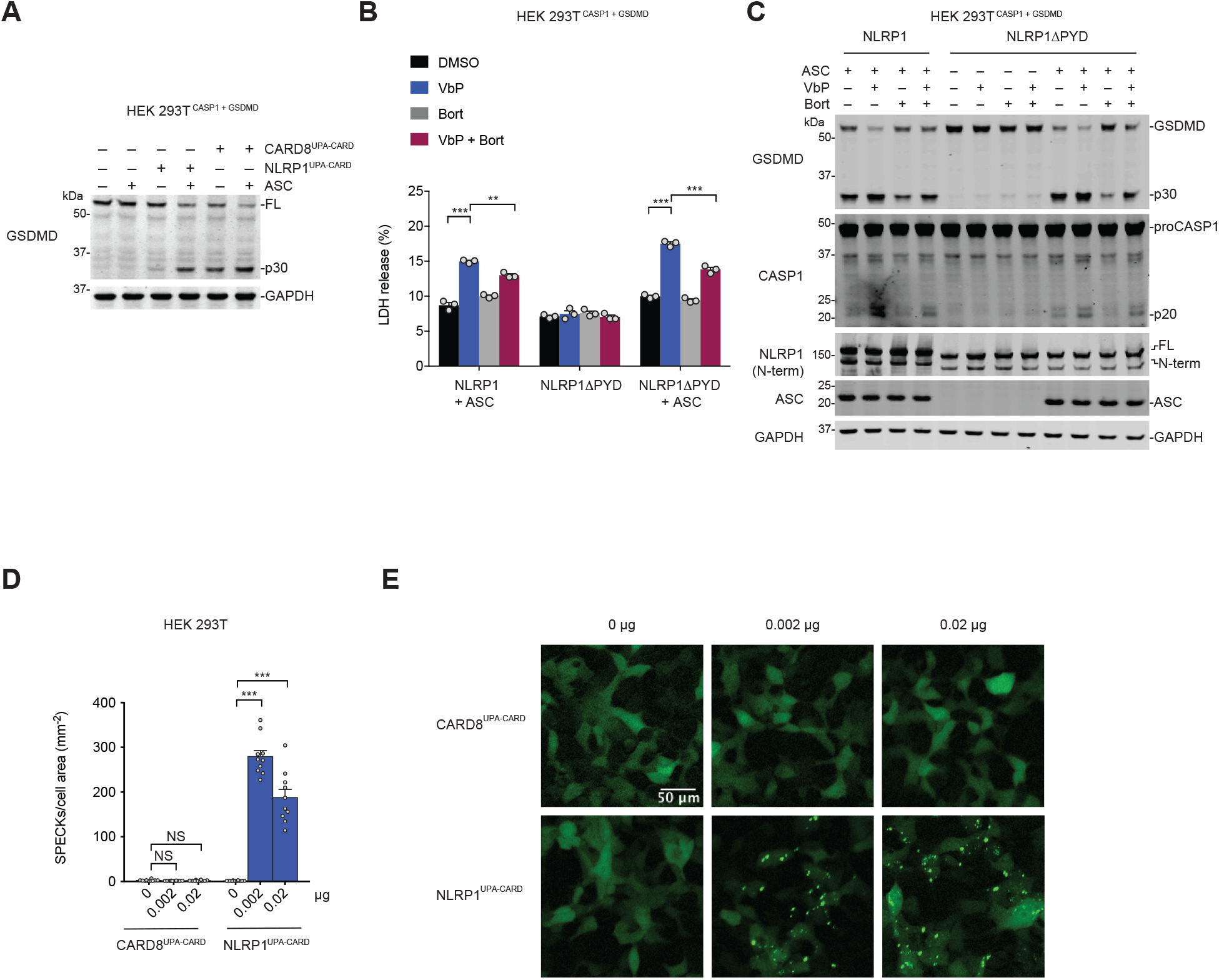
The NLRP1 CARD is responsible for inflammasome activation. **(A)** HEK 293T^CASP1 + GSDMD^ cells were transfected with plasmids encoding the UPA-CARD fragments of NLRP1 or CARD8 and ASC as indicated. After 24 h, lysates were evaluated by immunoblotting. **(B,C)** HEK 293T^CASP1 + GSDMD^ were transfected with plasmids encoding full-length NLRP1 or NLRP1 without a pyrin domain (NLRP1ΔPYD) and treated with DMSO or VbP (10 μM, 6 h). Supernatants were evaluated for LDH release **(B)** and lysates were analyzed by immunoblotting **(C)**. Data are means ± SEM of three biological replicates. ** *p* < 0.01, *** *p* < 0.001 by two-sided Students *t*-test. **(D,E)** HEK 293T cells were transfected with plasmids encoding GFP-tagged ASC and the UPA-CARD fragments of NLRP1 or CARD8, and then evaluated for ASC speck formation by fluorescence microscopy. Shown are the mean ± SEM **(D)** and representative images **(E)** from 10 technical replicates from one of two independent experiments. *** *p* < 0.001, by two-sided Students *t*-test. NS, not significant.

**Figure S2.**
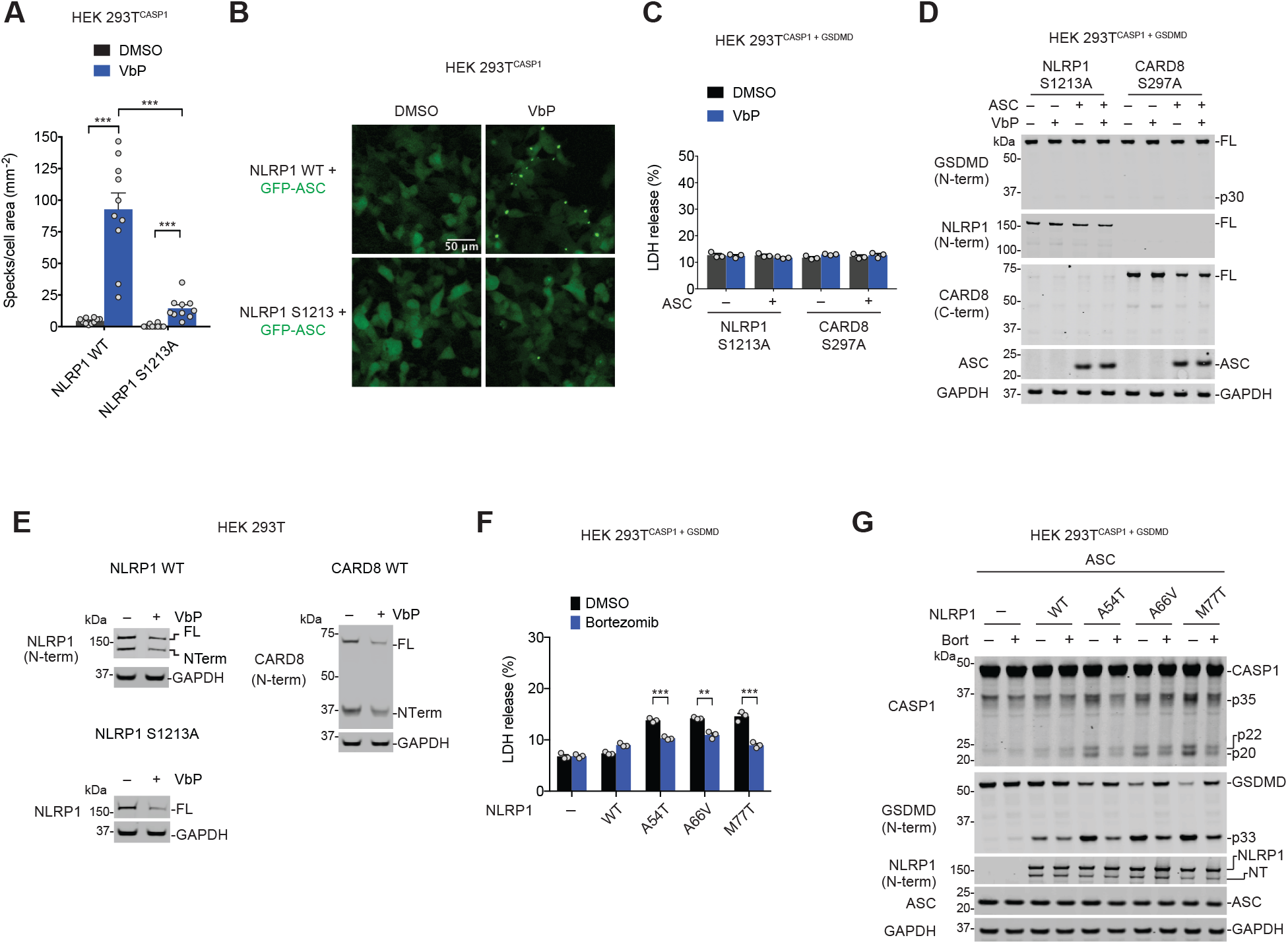
VbP and germline mutations activate NLRP1 via N-terminal degradation. **(A,B)** HEK 293T cells stably expressing CASP1 were transfected with plasmids encoding GFP-tagged ASC and the indicated NLRP1 protein, treated with DMSO or VbP (10 μM, 6 h), and evaluated for ASC speck formation by fluorescence microscopy. Shown are the mean ± SEM **(A)** and representative images **(B)** from 10 technical replicates from one of two independent experiments. *** *p* < 0.001, by two-sided Students *t*-test. **(C,D)** HEK 293T^CASP1 + GSDMD^ cells were transiently transfected with plasmids encoding autoproteolytic cleavage-deficient NLRP1 S1213A or CARD8 S297A with and without ASC, treated with VbP (10 μM, 6 h), and evaluated for LDH release **(C)** and GSDMD cleavage by immunoblotting **(D)**. The data are means ± SEM of three biological replicates. **(E)** HEK 293T cells were transiently transfected with the plasmids encoding CARD8 WT (0.1 μg), NLRP1 WT (0.5 μg), or NLRP1 S1213A (0.02 μg) prior to treatment with VbP (10 μM, 24 h). Cells were then treated with VbP again (10 μM, 24 h – total of 48 h of treatment) before lysates were evaluated by immunoblotting. **(F,G)** HEK 293T^CASP1 + GSDMD^ cells were transiently transfected with plasmids encoding the indicated NLRP1 protein and ASC for 24 h before being treated with bortezomib (20 μM, 6 h). Cell death was evaluated by LDH release **(F)** and lysates assessed by immunoblotting **(G)**. Data are means ± SEM of three biological replicates. ** *p* < 0.01, *** *p* < 0.001 by two-sided Students *t*-test.

**Figure S3.**
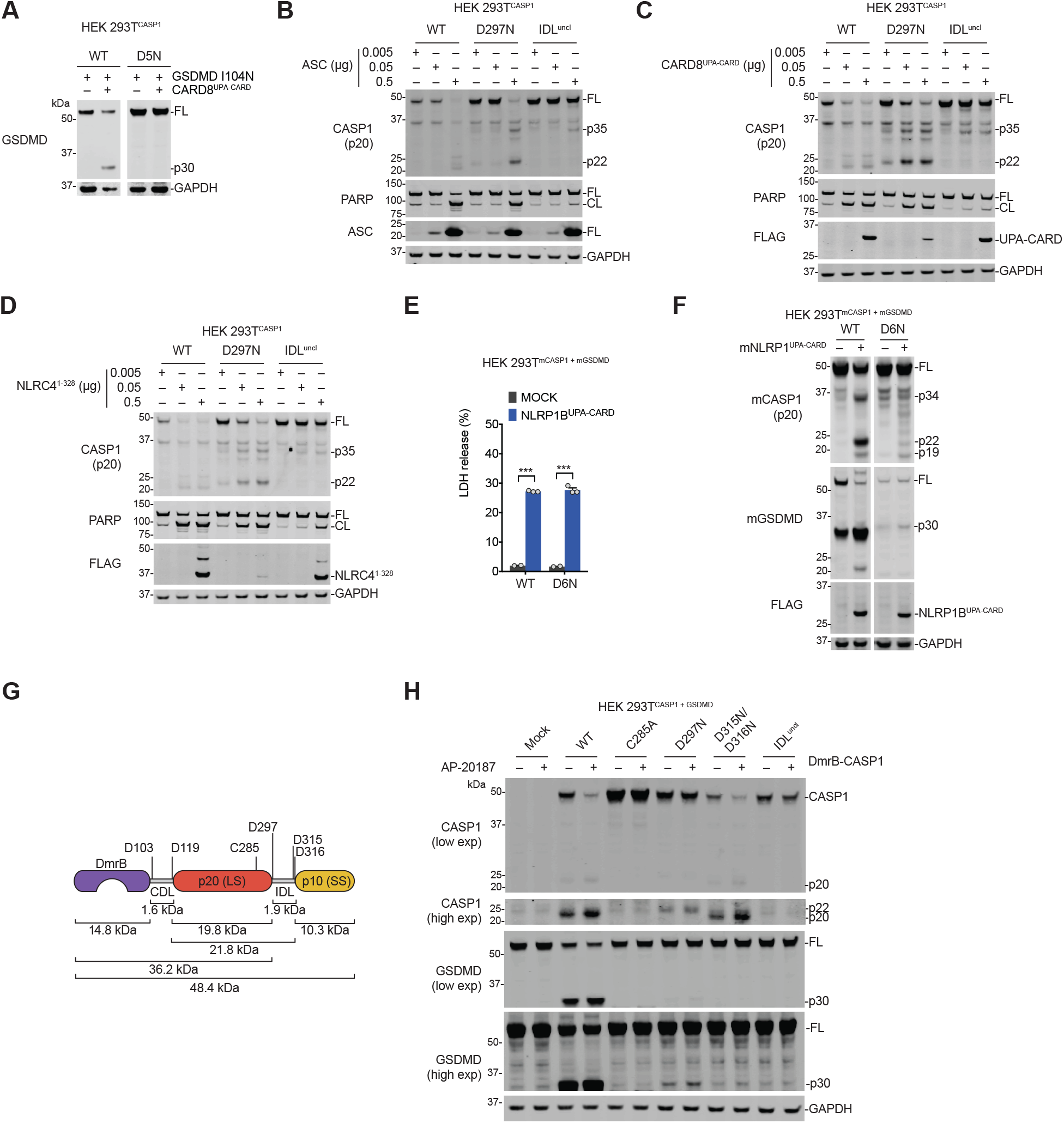
IDL cleavage is necessary for human pro-caspase-1 activation. **(A)** HEK 293T cells stably expressing CASP1 WT or CASP1 D5N were transiently transfected with plasmids encoding the UPA-CARD fragment of CARD8 (0.1 μg) and GSDMD I104N (0.1 μg). After 24 h, lysates were evaluated by immunoblotting. **(B-D)** HEK 293T cells stably expressing the indicated pro-caspase-1 were transiently transfected with the indicated amounts of plasmids encoding ASC **(B)** the UPA-CARD of CARD8 **(C)**, or residues 1-328 of NLRC4 **(D)**. Lysates were evaluated by immunoblotting after 24 h. **(E, F)** HEK 293T cells stably expressing mouse CASP1 WT or CASP1 D6N and mouse GSDMD were transiently transfected with a plasmid encoding the UPA-CARD fragment of NLRP1B (0.1 μg). After 24 h, supernatants were assessed for LDH release **(E)** and lysates were evaluated by immunoblotting **(F)**. Data are means ± SEM of two or three biological replicates. *** *p* < 0.001 by two-sided Students *t*-test. **(G)** Schematic of the DmrB-caspase-1 constructs. Predicted cleavage sites, sizes of potential cleavage products, and the catalytic cysteine are indicated. **(H)** HEK 293T cells stably expressing GSDMD were transiently transfected with the indicated DmrB-caspase-1 constructs for 24 h before addition of AP-20187 (500 nM, 1 h). GSDMD and CASP1 cleavage were evaluated by immunoblotting.

